# Viewing one’s body during encoding boosts episodic memory

**DOI:** 10.1101/318956

**Authors:** Lucie Bréchet, Robin Mange, Bruno Herbelin, Baptiste Gauthier, Andrea Serino, Olaf Blanke

## Abstract

Episodic autobiographical memories (EAMs) are recollections of contextually rich and personally relevant past events. EAM has been linked to the sense of self, allowing one to mentally travel back in subjective time and re-experience past events. However, the sense of self has recently been linked to online multisensory processing and bodily self-consciousness (BSC). It is currently unknown whether EAM depends on BSC mechanisms. Here, we used a new immersive virtual reality (VR) system that maintained the perceptual richness of life episodes and fully controlled the experimental stimuli during encoding and retrieval, including the participant’s body. We report that the present VR setup permits to measure recognition memory for complex and embodied 3D scenes during encoding and retrieval, that recognition memory depends on delay and number of changed elements, and that viewing one’s body as part of the virtual scene (as found in BSC studies) enhances delayed retrieval. This body effect was not observed when no virtual body or a moving control object was shown. These data show that embodied views improve recognition memory for 3D life-like scenes, thereby linking the sense of self, and BSC in particular, to episodic memory and the re-experiencing of specific past events in EAM.

## INTRODUCTION

A defining feature of episodic autobiographical memory (EAM) is the capacity to provide information about the content of our conscious personal experiences of “when” and “where” events occurred as well as “what” happened^1,2^. Previous studies defined EAM as the recall of contextually rich and personally relevant past events that are associated with specific sensory-perceptual and cognitive-emotional details ^3–10^. EAM has been distinguished from semantic autobiographical memory, the latter being associated with general self-knowledge and the recall of personal facts that are independent of re-experiencing specific past events ^11–17^.

In a series of seminal papers, Endel Tulving highlighted the subjective dimension of EAM associated with the re-experiencing of specific past events by pointing out the importance of the sense of self and introducing his influential notion of autonoetic consciousness. He argued that autonoetic consciousness is of fundamental relevance to EAM, allowing one to mentally travel back in subjective time and recollect one’s previous experiences^2,18–20^. Tulving distinguished autonoetic consciousness from noetic consciousness, linking the latter to semantic memory and semantic autobiographical memory and to knowing about (rather than re-experiencing) specific past events. Others extended Tulving’s notion of EAM and proposed that it is contributing to the sense of self across time ^10,12,21–25^ and developed behavioral tasks such as mental time travel ^26–31^.

Although, several other cognitive domains have been proposed to contribute to the sense of self (i.e. language, mental imagery, facial self-recognition^32–35^, recent research has highlighted the importance of non-cognitive multisensory and sensorimotor contributions to the sense of self. This novel theoretical and experimental approach is based on behavioral ^36,37^, neuroimaging ^38–40^ and clinical data ^39,41^ and involves the processing and integration of different bodily stimuli to the sense of self: bodily self-consciousness (BSC); for review see ^42,43^. BSC includes conscious experiences such as self-identification and self-location ^37,36,44,45^, as well as the first-person perspective ^39,46,47^. This work was based on clinical observations in neurological patients with so-called out-of-body experiences characterized by changes in the sense of self, in particular of the experienced self-location and perspective from an embodied first-person perspective to a third-person perspective ^39,41^ and has been able to induce milder, but comparable, states in healthy subjects using virtual reality (VR) technology to provide multisensory stimulation ^36,39,47^.

Given the link of BSC with subjective experience and previous claims that subjective re-experiencing of specific past events is a fundamental component of EAM ^2,18^, we argue that multisensory bodily processing may not only be of relevance for BSC, but also for consciousness concerning past events. Recent findings have shown that BSC impacts several perceptual and cognitive functions such as tactile perception ^48,49^, pain perception ^50,51^, visual perception ^52–54^, as well as egocentric cognitive processes ^55^. Concerning episodic memory, Bergouignan et al. ^56^ reported that recall of EAM items and hippocampal activity during the encoding of episodic events is modulated by the visual perspective from where the event was viewed during encoding and St. Jacques et al.^57^ showed that first-versus third-person perspective during retrieval modulated recall of autobiographical events and associated this with medial and lateral parietal activations. We here predicted that bodily multisensory processing that has been described to modulate BSC would interfere with EAM processes.

Traditionally, behavioral and neuroimaging EAM studies rely on questionnaires, verbal reports, interviews, or mental imagery and predominantly investigated memory retrieval by using a variety of stimuli and procedures such as cue words and pictures ^57–62^. For example, important research relied on interviews with the participants ^60,63^ on personalized lists of significant life events of participants ^9,30,64–66^, and employed different procedures asking participants to re-experience particular life episodes ^62,61,58,67,68^. This differs from research investigating verbal memory through encoding and recall of word lists ^69–72^ or testing spatial memory with figures, spatial paths, or other visuospatial materials ^73–75^. All these procedures, however, lack either the richness of real-life events or do not control the information during encoding. To overcome some of the limitations with respect to episodic memory, several EAM research groups have relied on advances in video technology and VR during encoding and retrieval of information (i.e. spatial navigation^76,77^; social interactions^78,79^). Participants were seated in front of a computer screen showing a virtual environment and asked to navigate in such environments using a joystick (encoding) and later asked to recall selected objects from the environment (retrieval). These computer-based VR studies suggest that both interactions with the environment during encoding or retrieval influence memory performance. Compared to passive participation, several VR studies showed better learning performances across free recall trials and recognition tasks ^76,80–82^. Plancher et al.^83^ suggested that interactions with the naturalistic environment created with VR enhanced spatial memory. However, despite these important achievements, these virtual environments were mostly using non-immersive VR systems, did not employ real life like virtual scenes, and did not use VR technology that allows integrating the participants’ body (and hence multisensory bodily stimulation) for the tested virtual life episodes. In the present experiments, we took advantage of a recently developed immersive VR system, which allows us to preserve the perceptual richness of life episodes, to fully control the experimental stimuli during encoding and retrieval, and to integrate and manipulate multisensory information of our participant’s body in an online fashion. The present experiments had one major technological and one major scientific goal: (1) develop and test EA-like memory in the laboratory with virtual episodes using immersive VR and (2) investigate whether multisensory bodily stimulations that have been shown to impact BSC, perception, and egocentric cognition modulates EAM.

In the first experiment, we tested our immersive VR system and sought to address some of the experimental limitations of earlier EAM studies, which either had limited control of actual autobiographical stimuli and events during encoding and only examined the stage of EAM retrieval ^5,59,66,84^ or controlled EAM encoding, but without the immersion into the original scenes during EAM retrieval ^9,56,64^. We further tested EAM performance and confidence for immersive three-dimensional (3D) VR scenes at two different time points and for different number of objects (that changed between both sessions), we predicted memory decreases depending on delay and on the number of objects changed. In the second experiment, we investigated the main hypothesis of the present experiments and tested the potential link between multisensory own body signals (that are fundamental for BSC) and EAM. We thus examined whether the presence of online and congruent multisensory cues from the subject’s body (i.e. the presence of one’s own physical body from the first-person viewpoint) impacts memory performance and confidence in the present VR paradigm, compared to an experimental condition where such online first-person bodily cues are absent. Finally, we performed a third (control) experiment in order to test whether the effect of multisensory bodily stimulation that we observed in the second experiment is specific to multisensory bodily cues.

## Methods

### Subjects

A total of 79 subjects with normal or corrected to normal vision were recruited to participate. None of the participants indicated neurological or psychiatric deficits. In experiment 1, 16 right-handed subjects (M = 23.7 years, SEM = 0.7 years, 8 female) participated in the immediate recognition group and 15 right-handed subjects (M = 23.4, SEM = 0.8, 7 female) participated in the one-hour delayed recognition group. In experiment 2, 16 right-handed subjects (M = 26.8 years, SEM = 0.6, 4 female) participated in the immediate recognition group and 16 right-handed subjects (M = 24.5 years, SEM = 1.1, 8 female) participated in the one-hour delayed recognition group. In experiment 3, 16 right-handed subjects (M = 25.4, SD = 3.7, 7 female) participated in one-hour delayed group. Sample size was derived from power analysis of previous studies^85,86^. Power analysis indicated that 16 participants are sufficient to perform a parametric analysis with a power of 0.8 (previous BSC studies). The study was approved by the local ethical committee and all three experiments were conducted in conformity with the Declaration of Helsinki. Informed consents were obtained from all our subjects.

### Virtual Reality Technology

Our VR technology uses a spherical capturing and recording system and an immersive setup for first-person perspective (1PP) replay of the recorded real environments. For recording a scene, 14 cameras (GoPro Hero4) are assembled on a spherical rig (360hero 3DH3PRO14H) and linked to 4 pairs of binaural microphones (3DIO Omni Binaural Microphone) to cover the entire sphere of perception around a viewpoint (360° horizontally and vertically, stereoscopic vision, binaural panoramic audio). A custom software (Reality Substitution Machine, RealiSM, http://lnco.epfl.ch/realism) then aggregates all data into a single high-resolution panoramic audiovisual computer format (equivalent to more than 4 stereoscopic full HD movies). A head-mounted display (HMD, Oculus Rift DK2; 900×1080 per eye, FOV ~105° Vertical, 95° Horizontal) was used to immerse subjects into the recording and sound was administered with noise-cancelling headphones (BOSE QC15). Furthermore, the HMD was coupled with a stereoscopic depth camera (Duo3D MLX, 752×480 at 56Hz) mounted on its front face to capture subjects’ bodies from 1PP. The RealiSM software then augments the fully immersive environment with a realistic view from which subjects could see their hands, trunk and legs from 1PP. As a result, subjects experienced as if they would be physically present in the pre recorded scenes and seeing oneself (not a 3D avatar). The software also allows integrating 3D virtual objects seamlessly in the scene (experiment 3).

### Stimuli

Subjects were immersed in three (experiment 1) or two (experiments 2 and 3) pre-recorded rooms via the HMD (see below). For the encoding session, 10 everyday-life objects (e.g., coffee machine, pen, trash bin) were placed in each room. These real-life objects created the natural context of episodic memory at both encoding and retrieval. During retrieval, rooms remained either exactly the same as during encoding (i.e. the same 10 real-life objects were again presented at the same places in the previously visited rooms) or some of the objects (i.e. 1, 2 or 3 objects) were replaced by new objects that were not previously seen in any of the scenes.

### Paradigm

Each of the three experiments consisted of two sessions, an incidental encoding period (session 1) followed by an immediate (group 1) or one-hour delayed (group 2) surprise recognition task (session 2). In all three experiments, subjects were not informed that we would later test their memory for the stimuli encountered during the encoding session. Before the two experimental sessions, subjects were seated on a chair and asked to put on the HMD and headphones. They could then familiarize themselves with the VR technology for several minutes. Paradigm and testing sequence are depicted in **Figure 1a**.

**Figure 1.**
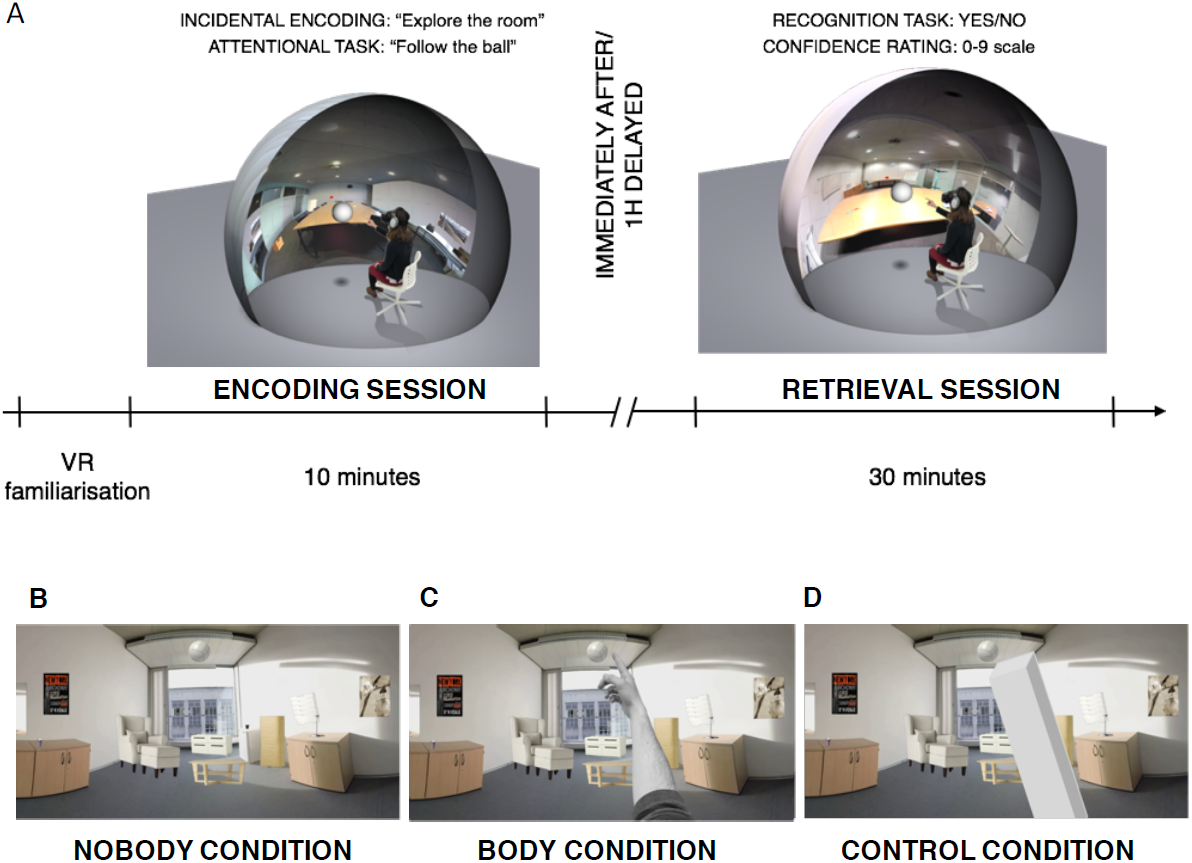
Experimental procedure and 3D scenes. After a period of familiarization with the immersive VR setup, participants performed the encoding session (10 minutes) during which they were exposed to different life-like 3D scenes (Figure 1A). Scenes were characterized by a room that contained different objects (table, photocopy machine, pen, etc.). In experiment 1, one group of participants performed the retrieval session (30 minutes) immediately after the encoding session or after a one hour delay (see main text for further detail). Figure 1B-D shows the different conditions during the encoding session that we used in experiments 1-3 (the retrieval session was the same across all experiments). Thus, participants always saw the same 3D scenes on the head-mounted display, but the body of the participant was either not seen at all (Figure 1B; no-body condition), seen as part of the 3D scene (Figure 1C; body condition), or instead of the body a control object was seen (Figure 1D; control condition).

### Encoding Session

During the encoding session, to assure that subjects explored the entire room and to monitor their attention within the different 3D scenes (i.e. the different rooms), participants were instructed to freely explore each virtual room. Moreover, we programmed a virtual ball that appeared in each of the three rooms and was moving within the rooms for a duration of 30 seconds and covered all sections of the virtual room. Participants were asked to fixate the virtual ball and to follow its movements through the virtual room. In total, the target object appeared at 6 different positions in each room. After the ball stopped moving, participants freely explored each room for another 30 seconds.

The procedure in experiment 2 was identical. However, in order to test the effect of viewing one’s own body during encoding we asked subjects to follow the trajectory of the ball and to point at the moving ball with their hand and finger. The main manipulation consisted in showing the participant’s physical body (body condition) or not (no-body condition). This was accomplished with the use of the stereoscopic depths cameras to capture in the participant’s body and by turning them on in the body and off in the no-body condition. The participant’s body was inserted in real-time in the virtual room and shown from the habitual visual first-person viewpoint. In the body condition, the subjects saw their physical hand, the trunk, and their legs (i.e. the stereoscopic depths cameras were turned on) in the HMD and as part of the virtual 3D scene **(Figure 1b).** In the no-body condition, the virtual 3D scene was identical except that the participant’s body was missing (i.e. the stereoscopic depths cameras were turned off) **(Figure 1c).** The order of presentation of the body and no-body condition was counterbalanced between subjects. In experiment 2 each participant explored two rooms (i.e. with 3 rooms as in experiment 1 the experiment would have been too long).

In experiment 3, participants were also asked to follow the movement of the ball appearing in each room (by physically pointing at it with their hand and finger). Yet in the object condition they were shown a non-bodily control object, instead of their own physical body **(Figure 1d**; see Supplemental video). The no-body condition was the same as in experiment 2. The presentation of the no-body and object condition was counterbalanced between subjects. No explicit instructions to memorize the objects of visited rooms were provided. In experiment 3, each participant explored two rooms (i.e. to keep conditions comparable with respect to experiment 2).

### Retrieval Session

During the retrieval session, which was the same for all three experiments (i.e. no body or control object was shown), subjects were informed that they would be immersed in the same rooms again. They performed a total of three blocks of 40 trials (each lasting 10 seconds). Within the three blocks of 40 trials, we presented 10 trials, which were exactly the same as during the original encoding session (i.e. including the same 10 objects). The remaining 30 trials were different and had either 1, 2 or 3 new objects replacing the respective number of objects shown during the encoding session. The blocks and individual trials in each block were presented in a randomized order. Participants were free to re-explore the virtual scenes for 10 seconds, after which they were asked two questions that were shown on the HMD. First, participants performed a two-alternative forced choice task (yes/no) whether the virtual scene shown during the retrieval session corresponded to the virtual scene during encoding (recognition task) (“Is the scene exactly the same as when you first saw it?”). Participants indicated their response with a wireless computer mouse. Second, participants were asked how confident they were about their answer (via a rating scale projected in the HDM; range from 0 (low) to 9 (high confidence)).

### Statistical analysis

In experiment 1, an independent samples t-test for hit rate and false alarm rate was applied to test whether ABM performance differed depending on delay (i.e. immediate x one-hour delayed condition). Independent sample t-test were further used to analyze whether the hit rate and false alarm for ABM confidence ratings differed depending on delay. A mixed analyses of variance (ANOVA) with the number of objects changed (i.e. 1 object, 2 objects or 3 objects) and delay (i.e. immediate x one-hour delayed group) was performed. Further, a 2 × 3 mixed ANOVA was run to understand the effects of delay (i.e. immediate x one-hour delayed groups) and number of objects changed in a room (i.e. 1 object, 2 objects, 3 objects) for the ABM confidence for the false alarm rates. Where appropriate, Greenhouse-Geisser corrections of degrees of freedom were used. Significant ANOVA effects were explored by post-hoc tests using Bonferroni correction. The significance level was set to alpha 0.05.

In experiment 2, we performed a mixed analysis of variance (ANOVA) with delay (i.e. immediate x one-hour delayed groups) and body (i.e. body x no-body condition) on ABM performance for hit rate and false alarm rate. Further, another 2 × 2 mixed ANOVA was performed for ABM confidence (for false alarm rates) with the factors retrieval time (i.e. immediate x one-hour delayed groups) and body (i.e. body present x body absent). Further, a 2 × 2 × 3 mixed analysis of variance (ANOVA) was performed in order to test the effects of delay (i.e. immediate x one-hour delayed groups), body (i.e. body x no-body condition) and the number of objects changed (i.e. 1 object, 2 objects or 3 objects). Similarly, a three-way mixed ANOVA was run to understand the effects of delay (i.e. immediate x one-hour delayed groups), the body condition (i.e. body x no-body condition) and the number of objects changed in a room (i.e. 1 object, 2 objects, 3 objects) on the ABM confidence (for the false alarm rates). Where appropriate, Greenhouse-Geisser corrections of degrees of freedom were used. Significant ANOVA effects were explored by post-hoc tests using Bonferroni correction. The significance level was set to alpha 0.05.

In experiment 3, an independent samples t-test was applied to test whether ABM performance differed in the no-body versus object condition. This was done for hit rate and for false alarm rate. An independent sample t-test was also used to examine whether ABM confidence false alarm differed in the no-body versus object condition. A mixed analyses of variance (ANOVA) with the number of objects changed (i.e. 1 object, 2 objects or 3 objects) and body (i.e. no-body x object) was performed. Similarly, a 2 × 3 mixed ANOVA was run to understand the effects of body (i.e. no-body x object) and number of objects changed in a room (i.e. 1 object, 2 objects, 3 objects) for the ABM confidence for the false alarm rates. Where appropriate, Greenhouse-Geisser corrections of degrees of freedom were used. Significant ANOVA effects were explored by post-hoc tests using Bonferroni correction. The significance level was set to alpha 0.05.

## Results

### Experiment 1 (Immediate versus one-hour delayed condition)

Participants in the delay group showed a significant decline in performance compared to the immediate memory recognition group. Mean hit rate was significantly lower in the delay group (M = 55.5, SEM = 5.3) than in the immediate group (M = 73.1, SEM = 3.6) (t (29) = 2.7, p = 0.01) **(Figure 2a).** False alarm rates did not differ between both groups (immediate group: M = 31.4, SEM = 5.8; delay group: M = 23.3, SEM = 3.0; t (29) = 1.1, p = 0.2) **(Figure 2b).** These data show that subjects recognized 3D scenic events better when tested immediately after the exposure than when tested with a delay of one hour, without any effect of delay on false recognitions.

**Figure 2.**
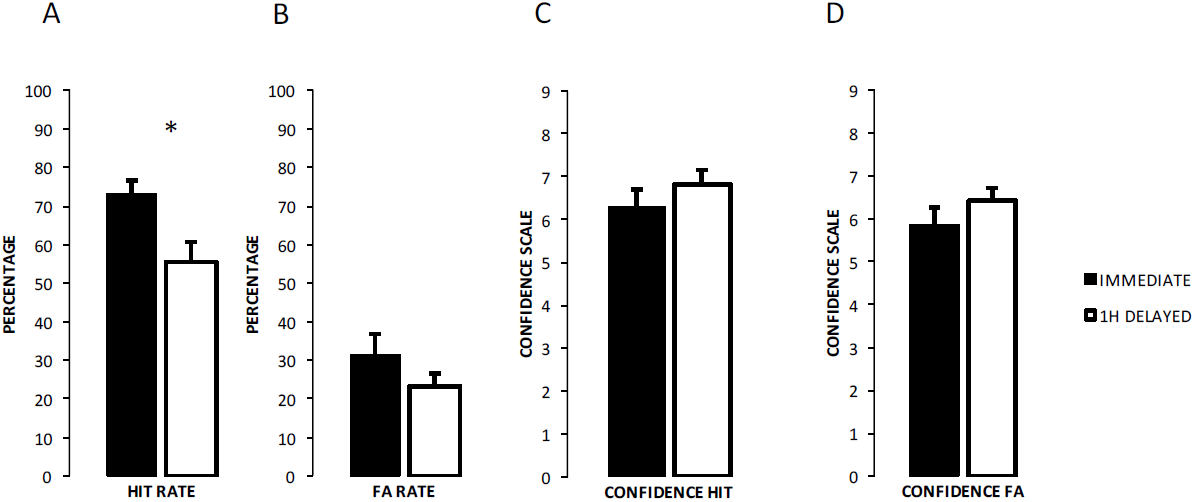
EAM performance in experiment 1 (immediate versus one-hour delay) EAM performance (hit rate, false alarm rates) and subjective confidence ratings are indicated in percentage + SEM. (**) P < 0.01; (*) P < 0.05. Figure 2A. Hit Rate; Figure 2B. False Alarm Rate; Figure 2C. Confidence ratings (Hits); Figure 2D. Confidence ratings (False alarms).

Confidence ratings for hits in the immediate group (M = 6.2, SEM = 1.6) were not significantly different from those in the delay group (M = 6.8, SEM = 1.3) (t (29) = 1.09, p = 0.2) **(Figure 2c)**. The same was found for false alarms confidence that did not differ between the immediate group (M = 5.8, SEM = 0.4) and delay group (M = 6.4, SEM = 1.2) (t (29) = 0.7, p = 0.3) **(Figure 2d).** Thus, despite changes in recognition, confidence did not differ depending on delay.

We next examined whether performance in the present task depended on the number of objects changed within each immersive 3D scene. This analysis was conducted on the false alarm rate (as no objects changed for hits, by definition). As predicted, analysis revealed a significant main effect for the number of objects changed (F (2, 58) = 52.85, p < 0.0005, partial η^2^ = 0.64) **(Figure 3a).** Pairwise comparisons were performed for statistically significant main effects and revealed that subjects made progressively fewer false alarms with increasing number of objects (all p-values < 0.0005).

**Figure 3.**
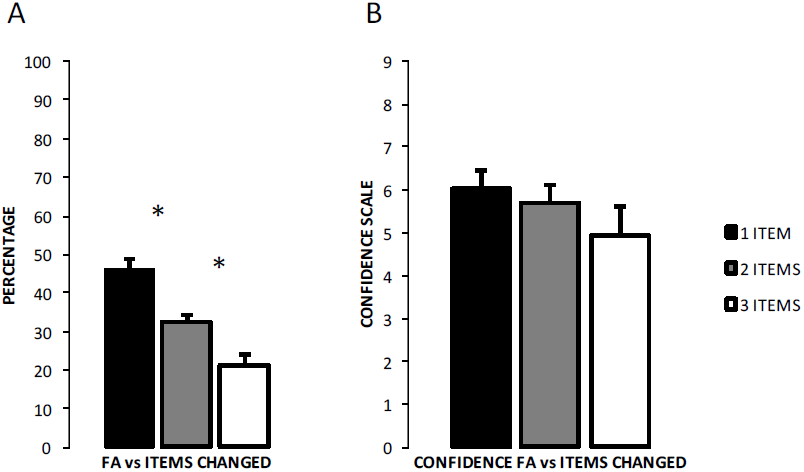
False alarms depend on number of items changed (experiment 1) EAM performance (false alarms) is indicated in percentage + SEM. (**) P < 0.01; (*) P < 0.05. Figure 3A. False Alarm versus Number of Items changed (i.e., 1 item, 2 items, 3 items); Figure 3B. Confidence Rate for False Alarm versus Number of Items changed (i.e., 1 item, 2 items, 3 items).

There was also a statistically significant main effect for the number of objects changed (F (2, 58) = 4.163, p = 0.02, partial η^2^ = 0.12) **(Figure 3b),** revealing that subjects were progressively more confident in their performance with increasing number of objects that were changed between both sessions. These data show that subjects made more recognition errors and were less confident in conditions in which less objects were changed between encoding and retrieval.

### Experiment 2 (Body versus no-body condition)

Data for hit rates showed a significant two-way interaction between the time of retrieval and body conditions (F (1,30) = 7.44, p = 0.01, partial η^2^ = .19). Post-hoc testing revealed that this effect was explained by a higher hit rate in the body, which was found specifically in the delay group (body: M = 82.5, SEM = 8.2; no-body condition: M = 63.7, SEM = 8.2; *t* (15) = 2.51, *p* = 0.02), but not in the immediate group (**Figure 4a**). The same analysis for false alarms rate did not reveal any differences F (1, 30) = 0.002, p =0.96, partial η^2^ = .00 (**Figure 4b**). These data show that recognition of immersive 3D scenes, that also include the first-person view of the subject’s body, mimicking real-life experience is modulated and enhanced with respect to the same scenes without such a bodily view.

**Figure 4:**
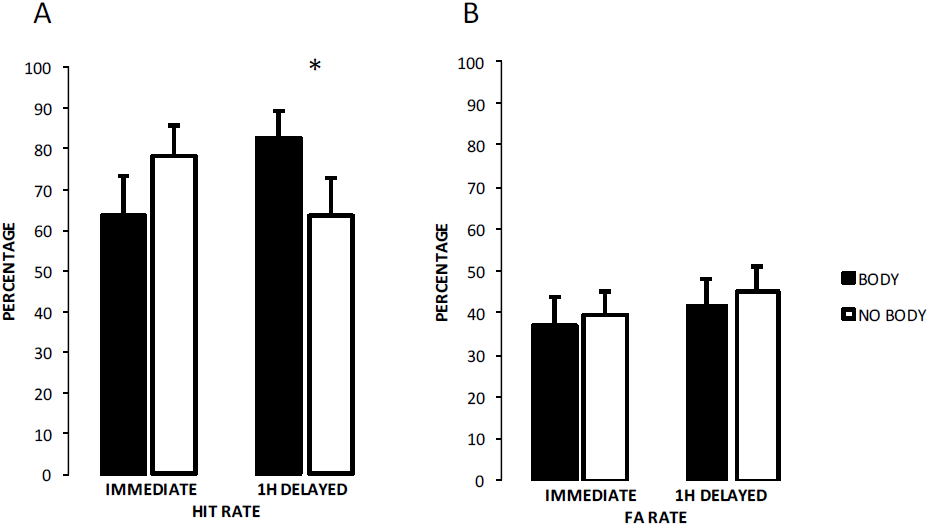
Body view enhances recognition (experiment 2) Immediate versus one-hour delay EAM performance is indicated in percentage + SEM is indicated. (**) P < 0.01; (*) P < 0.05. Figure 4A. Hit Rate in immediate versus 1h delayed; Figure 4B. False Alarm Rate in immediate versus 1h delayed.

Confidence ratings for hits did not reveal any differences between the time of retrieval and body conditions (F (1,30) = 1.06, p = 0.31, partial η^2^ = .03). The same analysis for false alarms also did not reveal any differences (F (1,30) = 0.193; p = 0.66, partial η^2^ = .00). Thus, despite changes in recognition, confidence did not differ depending on time of retrieval or body conditions.

We also examined whether memory performance in the immediate and delayed groups depended on number of objects changed and the body condition. The analysis revealed a significant main effect for the number of objects changed for the false alarm rate (F (2, 60) = 27.48, p < 0.0005, partial η^2^ = 0.47). Pairwise comparisons were performed for statistically significant main effect and revealed that subjects made progressively fewer false alarms with increasing number of objects (all p’s < 0.0005). No statistically significant three-way interaction was found between the time of retrieval, body conditions and number of objects changed (F (1.35, 40.54) = 1.84, *p* = 0.18, partial η^2^ = 0.05).

Similarly, we tested whether the confidence in the performance accuracy for both immediate and delayed groups (i.e. confidence ratings for false alarms trials) depended on the number of changed objects within each scene and the body condition. Results show that subjective ratings mirrored changes in memory performance. The main effect for objects showed a statistically significant difference for the number of objects changed (F (2, 60) = 7.79, p = 0.01, partial η^2^ =0.2). Post-hoc analysis revealed a statistically significant change from 1 object to 3 objects (p < 0.0005; Bonferroni corrected). There was no significant effect of body, nor significant three-way interaction was found (F (1.85, 55.55) = 1.14, *p* = 0.32, partial η^2^ = 0.03). Thus, despite changes in recognition of 3D scenes depending on whether the subjects viewed their body during encoding or not, our subjects’ confidence was equal across conditions. These data from experiment 2 show that subjects made more recognition errors and were less confident in conditions in which less objects were changed between encoding and retrieval, as in experiment1.

### Experiment 3 (object vs no-body condition)

There was no significant difference in hit rates for subjects in the object condition (M= 70.0, SEM = 8.3) compared to the no-body condition (M = 70.0, SEM = 8.2) (*t* (15) = 0.00, *p* = 1.00). Similarly, false alarm rates did not differ between groups (object group: M = 70.0, SEM = 8.3; no-body group: M = 70.0, SEM = 8.2; *t* (15) = 0.0, *p* = 1.0). These data show that recognition of immersive 3D scenes, where a non-bodily object, instead one’s own body, is visible from the first-person view, does not modulate performance in the present task with respect to the same scenes without body or rectangular control object.

Confidence for hits in the object condition (M = 4.8, SEM = 0.2) was not significantly different from the no-body condition (M = 4.4, SEM = 0.2). Confidence for false alarm also did not differ between conditions (object condition: M = 4.3, SEM = 0.3; no-body condition: M = 4.5, SEM = 0.2).

Further, we examined whether memory performance depended on number of objects changed and the body condition. The analysis revealed a significant main effect for the number of objects changed for the false alarm rate (F (2, 30) = 7.79, p < 0.0005, partial η^2^ = 0.34). Post-hoc analysis revealed a statistically significant change from 1 object to 3 objects (p = 0.01; Bonferroni corrected). No statistically significant two-way interaction was found between the body conditions and number of objects changed (F (2, 30) = 2.3, *p* = 0.11, partial η^2^ = 0.13). There was no significant difference between the no-body and object conditions.

We also tested whether the confidence in the performance accuracy depended on the number of changed objects within each scene and the body condition. The main effect for objects showed a statistically significant difference for the number of objects changed (F (2, 30) = 3.42, p = 0.04, partial η^2^ = 0.18). Similarly, no statistically significant two-way interaction was found between the confidence ratings for the body conditions and number of objects changed (F (2,30) = 0.55, *p* = 0.58, partial η^2^ = 0.03).

## DISCUSSION

The present study allows us to draw three major conclusions. First, the present VR setup permits to measure recognition memory for 3D scenes that are immersive, rich in contextual detail, and that further integrates the moving body of the participant in online fashion. Our VR setup, thus, approaches real-life experiences in controlled laboratory conditions. Moreover, the present VR setup allowed us to project the same 3D virtual scenes during the encoding and retrieval sessions, providing us arguably with a level of experimental control that is comparable to investigations in non-episodic memory. Second, applying this new setup we report that recognition memory for the tested VR scenes depends on the delay and on the number of changed elements between encoding and retrieval, comparable to findings for verbal and visual-spatial memory. Third, we show that viewing one’s body as part of the virtual scene during encoding enhances delayed retrieval. This body effect was not observed when no virtual body was shown or when a moving control object (instead of the virtual body) was shown, suggesting that embodied views lead to body-specific performance changes, as reported in studies investigating BSC.

### An experimental VR setup that controls real-life like episodes during encoding and retrieval

Most prior laboratory-based EAM studies used cue words or images to trigger memory retrieval and mental time travel to the past in a controlled fashion ^5,6,12,31,59,76,90,91,83,92^. However, these studies controlled only for memory retrieval but not for memory encoding^4^. Contrary to these previous studies, we exposed our participants to rich and immersive real-life scenes without the need for explicit mental time travel. Unlike earlier computer-based scenarios, we also did not present participants with artificial scenarios (simulated events in 3D), but immersed them into 360° video recordings of everyday real-life scenes that we digitalized for the encoding and retrieval sessions. Using the present naturalistic and controlled VR setup, we ensured that our participants experienced virtual 3D scenes with congruent multisensory bodily information (visual, motor, vestibular); these approach real-life experience as compared to classical virtual computer game tasks that have been used for episodic memory investigations in the past ^93,94^. Thus, the present VR technology and future improvements of it will open new possibilities for conducting episodic memory research under ecologically valid experimentation in the laboratory by providing not only the ability to precisely design all stimulus aspects, but also to replay fully controlled sequences of real-life events.

### Delay and number of changed objects modulates recognition memory performance

Our data reveal two classical episodic memory findings. Recognition memory for real-life like scenes decays with delay and improves depending on the number of items that were changed between encoding and retrieval. Previous EAM research is compatible with these findings, but has not been able to test or quantify this directly. Specifically, while associative recognition memory for words or pictures ^95–97^ and EAM ^98–100^ has been tested for different memory delays, previous VR-based paradigms, investigating the formation of episodic memory of lifelike events, mostly tested immediate memory performance ^77,78,101,102^ (but see ^103^). The present findings can be compared with classical memory findings for verbal and pictorial material where increasing delays increases forgetting ^18,104–107^ and with spatial memory work, where active navigation reduces forgetting as compared to passive viewing ^76,80,81,83^. Thus, although we only tested short delays (i.e. one hour), our data show that subjects remembered 3D scenes better when tested immediately after encoding as compared to delayed retrieval. Our second predicted finding that recognition memory was better when more items were changed between the encoding and retrieval is also compatible with classical findings concerning the recognition of visual changes when testing long-term memory for spatial scenes, complex figures (including faces), or short texts ^108,109^, further revealing the experimental validity of the present setup for research in episodic memory.

### Embodiment and episodic memory of life-like events

Besides reproducing classic memory effects, the present study also reveals a new finding, i.e. that memory is better when the body is visible at the encoding. Research on embodiment and BSC has used several VR paradigms and revealed the influence of multisensory and sensorimotor bodily input and has highlighted the importance of the view of the observer’s body ^43^. Such research showed that BSC can be modulated by showing the body or body parts of the participant from different first-person viewpoints compared to showing no body at all. Moreover, this effect has been shown to be body-specific by demonstrating that different non-corporeal objects shown from the same position and viewpoint do not alter BSC ^43^. Here, we extend this BSC principle to memory research by showing in experiment 2 that the recognition of 3D scenes that included within the first-person view also the subject’s body (as is characteristic of normal everyday perception) was modulated and significantly enhanced with respect to the same scenes without such a bodily view. This is compatible with previously reported effects for multisensory bodily perception ^48,49^ and BSC ^37,39,44^. These BSC studies showed that visuo-tactile perception, as well as self-identification and self-location towards a seen human body or body part are enhanced when the body is shown in congruent position with respect to the subject’s body. Accordingly, we argue that the present body effect on the recognition memory of 3D scenes is comparable to similar effects in multisensory perception and BSC (i.e. for review see ^43^) as well as a number of cognitive processes, where self-related bodily information is critical. For instance, viewing the body increases tactile perception ^110^, modulates interpersonal tactile responses ^111,112^, affects social cognition ^113,114^, and concept processing ^55^.

It could be argued that the enhanced EAM performance of experiment 2 could relate to differences in the amount of visual information provided in both conditions (higher in the body versus the no-body condition) or higher salience or attention due to the additional inclusion of the tracked body in the body condition. First, we note that addition of the tracked body actually covers or hides parts of the virtual scene and may have thus incidentally hid some of the changed items and should thus rather decrease recognition memory. Yet, the opposite was observed in experiment 2. However, in order to formally investigate the potential role of differences due to vision or attention between conditions we compared, in experiment 3, the no-body condition with a condition in which subjects viewed a non-bodily control object that was moving congruently with the participant’s body in real-time. Data from this experiment revealed no memory improvement in the object condition, arguing against a visual or attentional account and further corroborating our proposal that the present recognition enhancement is due to multisensory-motor bodily stimulation that has been shown to be crucial for BSC ^36,42,49,115^ and characteristic of normal everyday experience. These data also argue against the possibility that the present body effect on recognition memory can be generalized to an embodied object as the object condition did not induce any performance changes. By revealing bodily effects in the present EAM paradigm, we thus link BSC to EAM, extending earlier memory work ^56^ that has focused on contributions of the first-person perspective in autobiographical memory or of vestibular processing on EAM ^116^. Finally, based on these data we argue that the brain mechanisms of BSC are linked to those of autonoetic consciousness that are of fundamental relevance to EAM. Autonoetic consciousness is the ability to mentally travel back in subjective time and recollect one’s previous experiences ^2,18–20^ and the present data suggest that multisensory bodily processing during encoding and remembering are not only of relevance for the conscious bodily experiences of self-identification, self-location, and first-person perspective ^37,36,39,44–47^, but also autonoetic consciousness.

### Confidence and episodic memory

Does confidence mimic these changes in episodic memory performance? We report, as predicted, that confidence increased jointly with memory recognition improvements for conditions in which more objects were changed. This finding is in line with several studies showing that confidence in everyday, non-arousing EAM, measured by remember/know paradigms and recollection questionnaires, declines together with the objective memory performance ^95,98,99^. However, our data also show that confidence levels dissociate from memory performance, as delay dependency and the view of one’s body (experiment 2) during encoding modulated recognition memory, but not confidence levels. Further research needs to target objective memory performance and subjective confidence using real-life scenes as tested with the present VR setup. The differential delay- and body-effects in the present study suggest that memory performance and confidence rely on distinct functional mechanisms ^117^, potentially consistent with the classical two-component model of episodic memory highlighting the distinction between familiarity and recollection, with only the second leading to changes in confidence ^97^.

## References

1. Tulving, E. Episodic and semantic memory. (1972).

2. Tulving, E. Episodic Memory: From Mind to Brain. Annu. Rev. Psychol. (2002).

3. Svoboda, E., Mckinnon, M. C. & Levine, B. The functional neuroanatomy of autobiographical memoryl: A meta-analysis. Neuropsychologia 44, 2189–2208 (2006).

4. Cabeza, R. & St Jacques, P. Functional neuroimaging of autobiographical memory. Trends Cogn. Sci. 11, 219–27 (2007).

5. Daselaar, S. M. et al. The spatiotemporal dynamics of autobiographical memory: neural correlates of recall, emotional intensity, and reliving. Cereb. cortex 18, 217–29 (2008).

6. Oddo, S. et al. Specific role of medial prefrontal cortex in retrieving recent autobiographical memories: an fMRI study of young female subjects. Cortex 46, 29–39 (2010).

7. St Jacques, P. L., Kragel, P. a & Rubin, D. C. Dynamic neural networks supporting memory retrieval. Neuroimage 57, 608–16 (2011).

8. Addis, D. R. & Tippett, L. J. The contributions of autobiographical memory to the content and continuity of identity: A social-cognitive neuroscience approach. Self Contin. Individ. Collect. Perspect. 71–84 (2008).

9. St.Jacques, P. L., Montgomery, D. & Schacter, D. L. Modifying memory for a museum tour in older adults: Reactivation-related updating that enhances and distorts memory is reduced in ageing. Memory 23, 876–887 (2015).

10. Madore, K. P., Addis, D. R. & Schacter, D. L. Creativity and Memory. Psychol. Sci. 26, 1461–1468 (2015).

11. Tulving, E. What is episodic memory? (1993).

12. Levine, B. Autobiographical memory and the self in time: Brain lesion effects, functional neuroanatomy and lifespan development. Brain Cogn. 55 **55**, 54–68 (2004).

13. Rosenbaum, R. S. et al. Patterns of autobiographical memory loss in medial-temporal lobe amnesic patients. J. Cogn. Neurosci. 20, 1490–1506 (2008).

14. St-Laurent, M., Moscovitch, M., Tau, M. & Mcandrews, M. P. The temporal unraveling of autobiographical memory narratives in patients with temporal lobe epilepsy or excisions. Hippocampus 21, 409–421 (2011).

15. Kalenzaga, S. et al. Episodic memory and self-reference via semantic autobiographical memory: insights from an fMRI study in younger and older adults. Front. Behav. Neurosci. 8, 1–12 (2015).

16. Brown, A. D. et al. Episodic and semantic components of autobiographical memories and imagined future events in post-traumatic stress disorder. Memory 22, 595–604 (2014).

17. Chadwick, M. J. et al. Semantic representations in the temporal pole predict false memories. Proc. Natl. Acad. Sci. 113, 10180–10185 (2016).

18. Tulving. Memory and Consciousness. (1985). doi:10.1037/h0080017

19. Tulving, E. & Schacter, D. L. Priming and Human Memory Systems. Source Sci. New Ser. 247, 301–306 (1990).

20. Wheeler, M. A., Stuss, D. T. & Tulving, E. Towards a theory of episodic memory: the frontal lobes and automoetic consciousness. Psychol. Bull. 121, 331–354 (1997).

21. Conway, M. A. & Pleydell-pearce, C. W. The Construction of Autobiographical Memories in the Self-Memory System. Psychol. Rev. 107, 261–288 (2000).

22. Spreng, R. N. & Grady, C. L. Patterns of Brain Activity Supporting Autobiographical Memory, Prospection, and Theory of Mind, and Their Relationship to the Default Mode Network. J. Cogn. Neurosci. 22, 1112–1123 (2009).

23. Addis, Donna Rose and Tippett, L. The contributions of autobiographical memory to the content and continuity of identity: A social-cognitive neuroscience approach. Autobiographical Mem. identity 1–20 (2007).

24. Schacter, D. L. et al. The Future of Memoryl: Remembering, Imagining and the Brain. Neuron Rev. 76, 677–694 (2012).

25. Schacter, D. L., Benoit, R. G. & Szpunar, K. K. ScienceDirect Episodic future thinkingl: mechanisms and functions. Cobeha 17, 41–50 (2017).

26. Schacter, D. L., Addis, D. R. & Buckner, R. L. Remembering the past to imagine the future: the prospective brain. Nat. Rev. 8, 657–61 (2007).

27. Arzy, S., Thut, G., Mohr, C., Michel, C. M. & Blanke, O. Neural basis of embodiment: distinct contributions of temporoparietal junction and extrastriate body area. J. Neurosci. 26, 8074–81 (2006).

28. Arzy, S., Collette, S., Ionta, S., Fornari, E. & Blanke, O. Subjective mental time: the functional architecture of projecting the self to past and future. Eur. J. Neurosci. 30, 2009–17 (2009).

29. Arzy, S., Adi-Japha, E. & Blanke, O. The mental time line: an analogue of the mental number line in the mapping of life events. Conscious. Cogn. 18, 781–5 (2009).

30. Arzy, S., Molnar-Szakacs, I. & Blanke, O. Self in time: imagined self-location influences neural activity related to mental time travel. J. Neurosci. 28, 6502–7 (2008).

31. St Jacques, P. L., Olm, C. & Schacter, D. L. Neural mechanisms of reactivation-induced updating that enhance and distort memory. PNAS 110, 19671–8 (2013).

32. Araujo, H. F., Kaplan, J., Damasio, H. & Damasio, A. Neural correlates of different self domains. Brain Behav. 5, 1–5 (2015).

33. Gazzaniga, M. S., LeDoux, J. E. & Wilson, D. H. Language, praxis, and the right hemisphere: clues to some mechanisms of consciousness. Neurology 27, 1144–7 (1977).

34. Northoff, G. Self and brain: what is self-related processing? Trends Cogn. Sci. 15, 185–6 (2011).

35. Qin, P. & Northoff, G. How is our self related to midline regions and the default-mode network? Neuroimage 57, 1221–33 (2011).

36. Ehrsson, H. H. The Experimental Induction of Out-of-Body Experiences. Science (80-.). 317, 1048–1048 (2007).

37. Lenggenhager, B., Tadi, T., Metzinger, T. & Blanke, O. Video Ergo Suml: Manipulating Bodily Self Consciousness. Science (80-.). 1096–1099 (2007).

38. Guterstam, A. Decoding illusory self-location from activity in the human hippocampus. 9, 1–9 (2015).

39. Ionta, S., Gassert, R. & Blanke, O. Multi-sensory and sensorimotor foundation of bodily self-consciousness - an interdisciplinary approach. Front. Psychol. 2, 383 (2011).

40. Ionta, S., Martuzzi, R., Salomon, R. & Blanke, O. The brain network reflecting bodily self-consciousness: a functional connectivity study. Soc. Cogn. Affect. Neurosci. (2014). doi:10.1093/scan/nst185

41. Blanke, O., Landis, T., Spinelli, L. & Seeck, M. Out-of-body experience and autoscopy of neurological origin. Brain 127, 243–258 (2004).

42. Blanke, O. Multisensory brain mechanisms of bodily self-consciousness. Nat. Rev. Neurosci. 13, 556–71 (2012).

43. Blanke, O., Slater, M. & Serino, A. Behavioral, Neural, and Computational Principles of Bodily Self-Consciousness. Neuron 88, 145–166 (2015).

44. Petkova, V. I. & Ehrsson, H. H. If I were you: Perceptual illusion of body swapping. PLoS One 3, (2008).

45. Aspell, J. E. et al. Turning Body and Self Inside Out. Psychol. Sci. 24, 2445–2453 (2013).

46. Pfeiffer, C., Schmutz, V. & Blanke, O. Visuospatial viewpoint manipulation during full-body illusion modulates subjective first-person perspective. Exp. Brain Res. 232, 4021–4033 (2014).

47. Pfeiffer, C., Grivaz, P., Herbelin, B., Serino, A. & Blanke, O. Visual gravity contributes to subjective first-person perspective. Neurosci. Conscious. 2016, 1–12 (2016).

48. Aspell, J. E., Lenggenhager, B. & Blanke, O. Keeping in touch with one’s self: multisensory mechanisms of self-consciousness. PLoS One 4, e6488 (2009).

49. Noel, J. P., Pfeiffer, C., Blanke, O. & Serino, A. Peripersonal space as the space of the bodily self. Cognition 144, 49–57 (2015).

50. Hänsel, A., Lenggenhager, B., Von Känel, R., Curatolo, M. & Blanke, O. Seeing and identifying with a virtual body decreases pain perception. Eur. J. Pain 15, 874–879 (2011).

51. Romano, D., Llobera, J. & Blanke, O. Size and viewpoint of an embodied virtual body affect the processing of painful stimuli. J. Pain 17, 350–358 (2016).

52. Faivre, N., Arzi, A., Lunghi, C. & Salomon, R. Consciousness is more than meets the eye: a call for a multisensory study of subjective experience†. Neurosci. Conscious. 3, 1–8 (2017).

53. Salomon, R. et al. Unconscious integration of multisensory bodily inputs in the peripersonal space shapes bodily self-consciousness. Cognition 166, 174–183 (2017).

54. Salomon, R. et al. The Insula Mediates Access to Awareness of Visual Stimuli Presented Synchronously to the Heartbeat. J. Neurosci. 36, 5115–5127 (2016).

55. Canzoneri, E., di Pellegrino, G., Herbelin, B., Blanke, O. & Serino, A. Conceptual processing is referenced to the experienced location of the self, not to the location of the physical body. Cognition 154, 182–192 (2016).

56. Bergouignan, L., Nyberg, L. & Ehrsson, H. H. Out-of-body-induced hippocampal amnesia. PNAS 111, 4421–6 (2014).

57. St, P. L., Szpunar, K. K. & Schacter, D. L. Shifting visual perspective during retrieval shapes autobiographical memories. Neuroimage 148, 103–114 (2017).

58. Bluck, S. & Alea, N. Crafting the TALE: Construction of a measure to assess the functions of autobiographical remembering. Memory 19, 470–486 (2011).

59. Greenberg, D. L. et al. Co-activation of the amygdala, hippocampus and inferior frontal gyrus during autobiographical memory retrieval. Neuropsychologia 43, 659–674 (2005).

60. Kopelman, M. D., Wilson, B. A. & Baddeley, A. D. The autobiographical memory interview: a new assessment of autobiographical and personal semantic memory in amnesic patients. J. Clin. Exp. Neuropsychol. 11, 724–744 (1989).

61. Piolino, P., Desgranges, B. & Eustache, F. Episodic autobiographical memories over the course of time: cognitive, neuropsychological and neuroimaging findings. Neuropsychologia 47, 2314–29 (2009).

62. Rubin, D. C., Schrauf, R. W. & Greenberg, D. L. Belief and recollection of autobiographical memories. Mem. Cognit. 31, 887–901 (2003).

63. Levine, B., Svoboda, E., Hay, J. F., Winocur, G. & Moscovitch, M. Aging and autobiographical memory: Dissociating episodic from semantic retrieval. Psychol. Aging 17, 677–689 (2002).

64. Cabeza, R. et al. Brain activity during episodic retrieval of autobiographical and laboratory events: an fMRI study using a novel photo paradigm. J. Cogn. Neurosci. 16, 1583–94 (2004).

65. Svoboda, L. and. The effects of rehearsal on the functional neuroanatomy of episodic autobiographical and semantic remembering: an fMRI study. J. Neurosci. 29, 3073–3082 (2009).

66. Addis, D. R., Knapp, K., Roberts, R. P. & Schacter, D. L. Routes to the past: neural substrates of direct and generative autobiographical memory retrieval. Neuroimage 59, 2908–22 (2012).

67. Palombo, D. J., Williams, L. J., Abdi, H. & Levine, B. The survey of autobiographical memory (SAM): a novel measure of trait mnemonics in everyday life. Cortex 49, 1526–40 (2013).

68. Abram, M., Picard, L., Navarro, B. & Piolino, P. Mechanisms of remembering the past and imagining the future-New data from autobiographical memory tasks in a lifespan approach. Conscious. Cogn. 29, 76–89 (2014).

69. Kelley, W. M. et al. Hemispheric Specialization in Human Dorsal Frontal Cortex and Medial Temporal Lobe for Verbal and Nonverbal Memory Encoding. 20, 927–936 (1998).

70. Shallice, T. Brain Regions associated with acquisition and retrieval of verbal episodic memory. (1994).

71. Squire, L. R. Memory for relations in the short term and the long term after medial temporal lobe damage. Hippocampus 27, 608–612 (2017).

72. Tulving, E. Hemispheric encoding / retrieval asymmetry in episodic memoryl: Positron emission tomography findings. 91, 2016–2020 (1994).

73. Squire, L. R. Nondeclarative Memory: Multiple Brain Systems Supporting Learning. J. Cogn. Neurosci. 4, 232–243 (1992).

74. Moscovitch, M. et al. Functional neuroanatomy of remote episodic, semantic and spatial memory: a unified account based on multiple trace theory. J. Anat. 207, 35–66 (2005).

75. Eichenbaum, H. & Cohen, N. J. Can We Reconcile the Declarative Memory and Spatial Navigation Views on Hippocampal Function? Neuron 83, 764–770 (2014).

76. Sauzéon, H. et al. The use of virtual reality for episodic memory assessment: effects of active navigation. Exp. Psychol. 59, 99–108 (2011).

77. Schedlbauer, A. M., Copara, M. S., Watrous, A. J. & Ekstrom, A. D. Multiple interacting brain areas underlie successful spatiotemporal memory retrieval in humans. Sci. Rep. 4, 6431 (2014).

78. Burgess, N., Maguire, E. a, Spiers, H. J. & O’Keefe, J. A temporoparietal and prefrontal network for retrieving the spatial context of lifelike events. Neuroimage 14, 439–53 (2001).

79. Hawco, C. et al. Source retrieval is not properly differentiated from object retrieval in early schizophrenia: An fMRI study using virtual reality. NeuroImage Clin. (2014). doi:10.1016/j.nicl.2014.08.006

80. James, K. H., Humphrey, G. K. & Goodale, M. a. Manipulating and recognizing virtual objects: where the action is. Can. J. Exp. Psychol. 55, 111–120 (2001).

81. James, K. H. et al. ‘Active’ and ‘passive’ learning of three-dimensional object structure within an immersive virtual reality environment. Behav Res Methods Instrum Comput 34, 383–390 (2002).

82. Hahm, J. et al. Effects of active navigation on object recognition in virtual environments. Cyberpsychol. Behav. 10, 305–308 (2007).

83. Plancher, G., Barra, J., Orriols, E. & Piolino, P. The in fluence of action on episodic memory: A virtual reality study. Q. J. Exp. Psychol. 1–15 (2012).

84. Bonnici, H. M. et al. Detecting representations of recent and remote autobiographical memories in vmPFC and hippocampus. J. Neurosci. 32, 16982–91 (2012).

85. Park, H.-D. et al. Transient Modulations of Neural Responses to Heartbeats Covary with Bodily Self-Consciousness. J. Neurosci. 36, 8453–8460 (2016).

86. Blanke, O. et al. Neurological and robot-controlled induction of an apparition. Curr. Biol. 24, 2681–2686 (2014).

87. Green, D.M. and Sweets, J. A. Signal Detection Theory and Psychophysics. (1966).

88. Maniscalco, B. & Lau, H. A signal detection theoretic approach for estimating metacognitive sensitivity from confidence ratings. Conscious. Cogn. 21, 422–430 (2012).

89. Morey, R. D., Rouder, J. N. & Jamil, T. Computation of Bayes Factors for Common Designs. 54 (2015).

90. Addis, D. R., Moscovitch, M., Crawley, A. P. & McAndrews, M. P. Recollective qualities modulate hippocampal activation during autobiographical memory retrieval. Hippocampus 14, 752–62 (2004).

91. Holland, A. C., Addis, D. R. & Kensinger, E. a. The neural correlates of specific versus general autobiographical memory construction and elaboration. Neuropsychologia 49, 3164–77 (2011).

92. Buchy, L. et al. Functional magnetic resonance imaging study of external source memory and its relation to cognitive insight in non-clinical subjects. Psychiatry Clin. Neurosci. 68, 683–91 (2014).

93. Burgess, N., Becker, S., King, J. a & O’Keefe, J. Memory for events and their spatial context: models and experiments. Philos. Trans. R. Soc. Lond. B. Biol. Sci. 356, 1493–503 (2001).

94. Ekstrom, A. D., Copara, M. S., Isham, E. a, Wang, W. & Yonelinas, A. P. Dissociable networks involved in spatial and temporal order source retrieval. Neuroimage 56, 1803–13 (2011).

95. Dolcos, F., LaBar, K. S. & Cabeza, R. Remembering one year later: role of the amygdala and the medial temporal lobe memory system in retrieving emotional memories. PNAS 102, 2626–2631 (2005).

96. Tompary, A., Duncan, K. & Davachi, L. Consolidation of Associative and Item Memory Is Related to Post-Encoding Functional Connectivity between the Ventral Tegmental Area and Different Medial Temporal Lobe Subregions during an Unrelated Task. J. Neurosci. 35, 7326–31 (2015).

97. Yonelinas, A. P. Components of episodic memory: the contribution of recollection and familiarity. Philos. Trans. R. Soc. B Biol. Sci. 356, 1363–1374 (2001).

98. Talarico, J. M., Rubin, D. C., King, M. L. & Brown, J. Flashbulb memories. Psychol. Sci. 14, 455–461 (2003).

99. Hirst, W. & Phelps, E. A. Flashbulb Memories. Curr. Dir. Psychol. Sci. 25, 36–41 (2016).

100. Stone, C. B., Luminet, O. & Hirst, W. Induced forgetting and reduced confidence in our personal past? The consequences of selectively retrieving emotional autobiographical memories. Acta Psychol. (Amst). 144, 250–257 (2013).

101. Spiers, H. J. et al. Unilateral temporal lobectomy patients show lateralized topographical and episodic memory deficits in a virtual town. Brain 124, 2476–2489 (2001).

102. Watrous, A. J., Tandon, N., Conner, C. R., Pieters, T. & Ekstrom, A. D. Frequency-specific network connectivity increases underlie accurate spatiotemporal memory retrieval. Nat. Neurosci. 16, 349–56 (2013).

103. Plancher, G., Tirard, A., Gyselinck, V., Nicolas, S. & Piolino, P. Using virtual reality to characterize episodic memory profiles in amnestic mild cognitive impairment and Alzheimer’s diseasel: Influence of active and passive encoding. Neuropsychologia 50, 592–602 (2012).

104. Sharot, T. & Phelps, E. a. How arousal modulates memory: disentangling the effects of attention and retention. Cogn. Affect. Behav. Neurosci. 4, 294–306 (2004).

105. Dolcos, F., LaBar, K. S. & Cabeza, R. Interaction between the amygdala and the medial temporal lobe memory system predicts better memory for emotional events. Neuron 42, 855–863 (2004).

106. Sharot, T., Martorella, E. a, Delgado, M. R. & Phelps, E. a. How personal experience modulates the neural circuitry of memories of September 11. PNAS 104, 389–94 (2007).

107. Talamini, L. M. & Gorree, E. Aging memories: Differential decay of episodic memory components. Learn. Mem. 19, 239–246 (2012).

108. Landauer, T. How much do people remember? Some estimates of the quantity of information in long term memory. Cogn. Sci. 10, 477–493 (1986).

109. Shepard, R. N. Recognition memory for words, sentences, and pictures. J. Verbal Learning Verbal Behav. 6, 156–163 (1967).

110. Serino, A. & Haggard, P. Touch and the body. Neurosci. Biobehav. Rev. 34, 224–236 (2010).

111. Serino, A., Pizzoferrato, F. & Làdavas, E. Viewing a Face (specially one’s own face) being touched enhances tactile perception on the face. Psychol. Sci. 19, 434–438 (2008).

112. Keysers, C. et al. A touching sight: SII/PV activation during the observation and experience of touch. Neuron 42, 335–346 (2004).

113. Maister, L., Slater, M., Sanchez-Vives, M. V. & Tsakiris, M. Changing bodies changes minds: Owning another body affects social cognition. Trends Cogn. Sci. 19, 6–12 (2015).

114. Keysers, C., Kaas, J. H. & Gazzola, V. Somatosensation in social perception. Nat. Rev. Neurosci. 11, 726–726 (2010).

115. Serino, A. et al. Bodily ownership and self-location: Components of bodily self-consciousness. Conscious. Cogn. 22, 1239–1252 (2013).

116. Dijkstra, K., Kaschak, M. P. & Zwaan, R. A. Body posture facilitates retrieval of autobiographical memories. Cognition 102, 139–149 (2007).

117. Sadeh, T., Ozubko, J.D., Winocur, G., & Moscovitch, M. How we forget may depend on how we remember. Trends Cogn. Sci. (2014).

